# Unambiguous observation of vesicle fusion in the interior of a cone

**DOI:** 10.1101/2020.06.20.162636

**Authors:** Ge Zhenpeng

## Abstract

When a vesicle was put onto the inner surface of a cone, it would spontaneously move to the bottom of the cone. The mechanism was explained elsewhere. In this work, we showed that when we put two vesicles onto the inner surface of a cone, both of them would spontaneously climb to the bottom of the cone. What’s following is the unambiguous fusion of these two vesicles. The probability of fusion was greatly enhanced by the confined space and close contact between these two vesicles. Our result may shed new insights on the fusions of vesicles in complex environments in biological systems, especially in the axonal and nerve growth processes.

## Introduction

Vesicle fusion is the merging between two vesicles, or more generally between a vesicle and a part of a cell membrane or other target cell compartments, such as a lysosome. For example, in exocytosis^1-3^, vesicle fusion is the end stage of secretion from secretory vesicles, where their contents are expelled from the cell.

The close contact between two vesicles is often the prerequisite of vesicle fusion^4^. However, due to the electrostatic interaction and hydration shells, vesicles usually repulse each other. To achieve close contact between vesicles, SNARE proteins^5-7^ as well as calcium^8-10^ often play import roles.

In this work, we showed that by carefully designing a conical structure, close contact between vesicles was ensured and the fusion was greatly enhanced by the close contact between vesicles. In particular, in our toy model we put two vesicles onto the inner surface of a cone. The two vesicles spontaneously moved to the bottom of cone. The driving force is attributed to the demand of maximizing the adhesion energy with the substrate while maintaining its structure, which was explained in detail in our previous work.^11^ After both reaching the bottom of the cone, the two vesicles were “forced” to contact each other and finally fuse with each other after long-term contact and gradually changing poses. Our result may shed new insights on the fusions of vesicles in complex environments in biological systems, especially in the axonal and nerve growth processes.

## Methods

The solvent-free force field from Cooke^12,13^ was adopted for vesicles in this work. In this model, each lipid molecular contains one head bead and two tail beads. The interactions between the beads are governed by a combination of Weeks-Chandler-Andersen potential^14^, attractive potential as well as additional bonding and bending terms. The unit simulation timescale is τ ≈ 10 ns, which is estimated upon lipid self-diffusion simulations. The unit length is σ ≈ 1 nm, which is estimated by comparing the thickness of a bilayer from experiments and simulations.

The substrate is built by first constructing a cube with the face-centered cubic structure and then cutting it into a conical shell. The half-angle is set to be 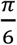. The length between adjacent beads in the cube is set to be 1σ. The interaction between the substrate and vesicle is modeled as the same with that of the tail-tail interaction in vesicles, which has the following form.

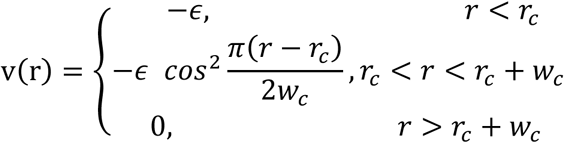

The diameter of the vesicle in this study is set to be 6 nm, which ensures its stability in solution.

**Fig 1.**
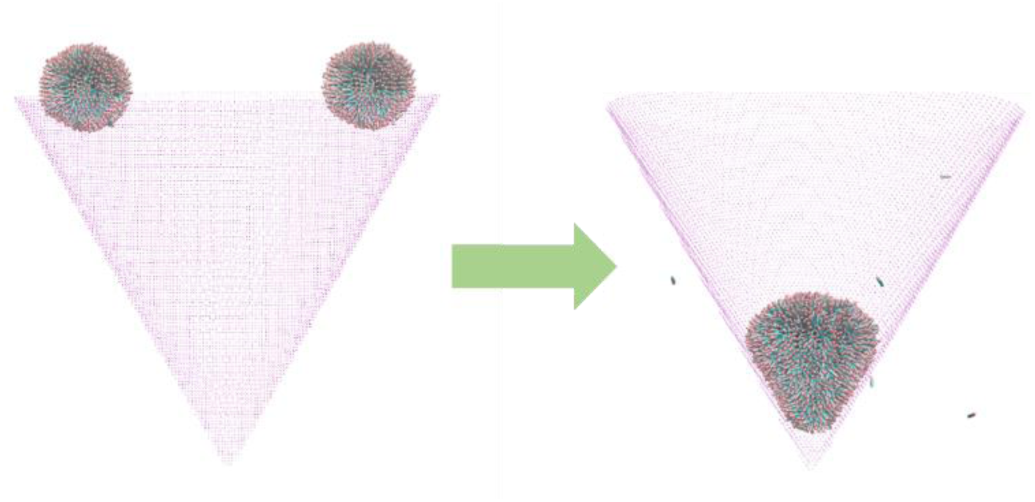
A schematic representation of the fusion between two vesicles in the interior of a cone.

## Results and discussions

The total time of our simulation is 14550 τ. At the beginning of the simulation, both of the vesicles adhered to the cone indicating a favorable interaction between the vesicles and cone. The two vesicles then adjusted its shape on the cone to maximize the adhesion energy for a short time. After that, the vesicle on the right began to move directionally towards the bottom of the cone. At 5370 τ, it reached the bottom and started to maximize its contact with the cone.

**Fig 2.**
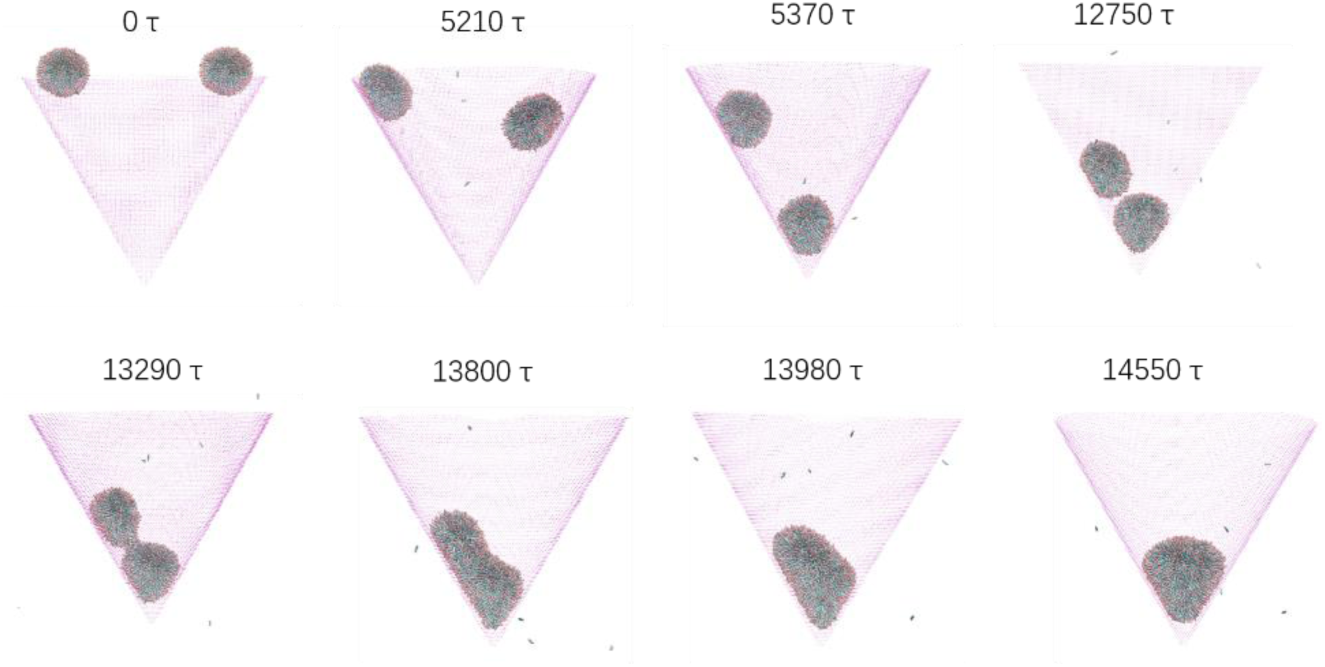
Snapshots extracted from the simulation showing the directional motion and subsequent fusion of the vesicles.

The vesicle on the left moved a little bit slower. Since the bottom had already been occupied by the other vesicle, it could only reach the upper position of the other vesicle. Then the two vesicles began to contact, but the contact is not stable. They continuously contact and left each other and meanwhile adjusted their shapes to ensure a better contact. This process lasted for a pretty long time. The turning point emerged when a bridge between the two vesicles appeared. After time, the two vesicles merged each other quickly and then became a large vesicle. The new vesicle then “rotated” its head to an axisymmetric shape. We also plotted the z coordinates of the center of the vesicles in Fig 3, which has a clear depiction of the whole process.

**Fig 3.**
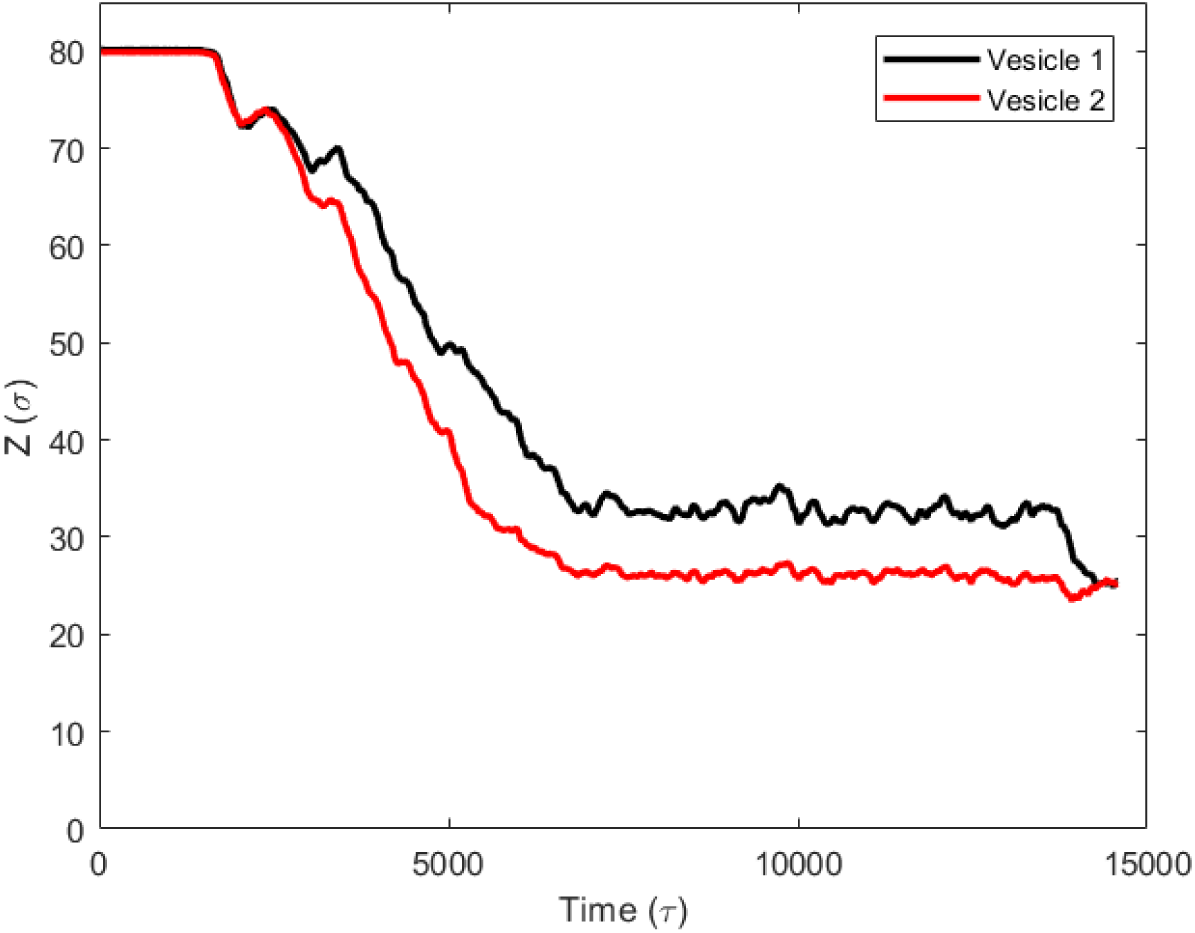
The z coordinates of the center of the vesicles.

## Conclusion

In this work, we observed the Unambiguous fusion between two vesicles on the inner surface of a cone. Due to the demand of maximizing its adhesion energy with the substrate, vesicles spontaneously moved to the bottom of the cone. Then a close contact between the vesicles was ensured, which facilitated the subsequent fusion in the confined space. The fusion was achieved by the continuous adjustment of the shapes of the vesicles with a bridge being the key medium.

## Supporting information

Vesicle fusion in a cone

